# OrganellarGenomeDRAW (OGDRAW) version 1.3.1: expanded toolkit for the graphical visualization of organellar genomes

**DOI:** 10.1101/545509

**Authors:** Stephan Greiner, Pascal Lehwark, Ralph Bock

## Abstract

- OGDRAW has become the standard tool for displaying maps of organellar genomes
- it converts GenBank entries into graphical maps
- a new version with improved functionality has been released

**Abstract:** Organellar (plastid and mitochondrial) genomes play an important role in resolving phylogenetic relationships, and next-generation sequencing technologies have led to a burst in their availability. The ongoing massive sequencing efforts require software tools for routine assembly and annotation of organellar genomes as well as their display as physical maps. OrganellarGenomeDRAW (OGDRAW) has become the standard tool to draw graphical maps of plastid and mitochondrial genomes. Here were present a new version of OGDRAW equipped with a new front end. Besides several new features, OGDRAW has now access to a local copy of the organelle genome database of the NCBI RefSeq project. Together with batch processing of (multi-)GenBank files, this enables the user to easily visualize large sets of organellar genomes spanning entire taxonomic clades. The new OGDRAW server can be accessed at https://chlorobox.mpimp-golm.mpg.de/OGDraw.html.

## Introduction

Organellar genomes display relatively conserved gene contents, are usually transmitted uniparentally (most often maternally) and are, therefore, excluded from sexual recombination. In most taxonomic groups, the mitochondrial and plastid genomes are small and occur in many copies per cell, which makes them convenient (and cheap) targets of sequencing projects (1-3). Moreover, these properties make organellar genomes extremely informative in resolving taxonomic relationships, and it can be expected that the enormous increase in published organellar genome sequences will continue for the foreseeable future (2,3). As next-generation sequencing technologies led to a massive increase in available sequence information, they also pushed forward technology development in the area of organellar genome assembly and annotation (1,3). There are many assembly methods and pipelines [e.g., GetOrganelle or IOGA; (4,5)], and we are currently aware of no less than twelve (semi-)automatic annotation tools for organellar genomes (1,3,6-15). These range from specialized applications such as MITOS (11), that were designed for a subset of organellar genomes and whose output requires little to no quality control or manual curation, to GeSeq, a flexible tool that allows the annotation of essentially any organellar genome (15). In addition, command line tools that can be incorporated into assembly pipelines such as Plann (13) are available, as well as annotation tools that have implemented downstream data processing (e.g., for phylogenetic analyses), as for example, Verdant (14).

The large diversity of sequencing and annotation software for organellar genomes is in stark contrast to the very small number of tools suitable to visualize finalized genome records. With the exception of metazoan mitochondrial genomes (16), most organellar genomes are too large for standard plasmid drawing programs and hence difficult to display graphically. Before the launch of OrganellarGenomeDRAW (OGDRAW) in 2007 (17), organelle genome maps were often drawn manually, lacked a homogeneous design and were inconsistent in feature display. These shortcomings were overcome by OGDRAW, which quickly became the standard in the field and currently, is the only tool that is widely used to generate graphical maps of organellar genomes. As of January 2019, OGDRAW has received more than 1,000 citations (Google Scholar), and the OGDRAW server generates about 120 maps per day. When the underlying operation system of the original OGDRAW server became outdated (CentOS v6.7, end of life May 2017), we decide to incorporate OGDRAW into software toolbox CHLOROBOX (https://chlorobox.mpimp-golm.mpg.de) that has been developed at the Max Planck Institute of Molecular Plant Physiology in Potsdam-Golm (MPI-MP). CHLOROBOX offers software applications for the analyses of (mainly plant-derived) nucleic acid and protein sequences. As described in detail below, in the course of the movement of OGDRAW to its new environment, we added several new features to the program (in its new version 1.3.1) and fixed known bugs.

## Result and Discussion

### Program description

OGDRAW converts annotations in GenBank format to graphical maps. The input file must be a GenBank flat file (https://www.ncbi.nlm.nih.gov/Sitemap/samplerecord.html), whereas the output file can be generated in different file formats. OGDRAW can produce bitmap or vector graphics at a range of resolutions (bitmap). Maps of both circular and linear genomes can be drawn. Coding regions and other feature-bearing regions of organellar genomes (and other DNA molecules such as plasmids) are visualized, and gene expression data can be displayed. The program can also display cut sites of restriction enzymes. For details on the basic functionality and implementation of OGDRAW, the interested reader is referred to the publications describing OGDRAW v1.0 in 2007 (17) and v1.2 in 2013 (18).

### The new front end

We equipped OGDRAW with a new front end that, from a technical perspective, greatly increases its user friendliness. As the previous version was a server-side rendered implementation, the user had to wade through several pages, including file upload, parameter input and result download. Comparing different input parameters was a cumbersome task, since for every job, the whole process needed to be restarted. By contrast, the new version of OGDRAW represents a state-of-the-art single-page application (SPA) with asynchronous client-server communication. Among other useful features, the site now also “remembers” previously set parameters. This facilitates a much easier comparison of outputs, for example, when comparing maps of the same organellar genomes with different resolutions.

### The new GUI

The graphical user interface (GUI) of OGDRAW v1.3.1 was adjusted to the CHLOROBOX design and consists of three columns (Figures 1 and 2). In the first column (I), the user can select the mode (standard or transcript; box Ia) and upload the required GenBank files. Genome conformation (circular or linear) and sequence source (plastid, mitochondrial, or other) are automatically extracted from the GenBank entry (box Ib). Here, the user can also select the “tidy up” option. Upon selecting this option, OGDRAW to ignore very long gene names that likely represent annotation errors. In addition, it will reformat many gene names that do not meet nomenclature conventions [for details, see refs. (17,18)]. In the second column (II), depending on the mode chosen, the user can upload either a custom configuration as XML file (Figure 1, box IIa) or gene expression datasets (Figure 2, box IIa). Customizing a configuration file offers the possibility to display features that are not included in the standard configuration of OGDRAW such as CDS, mRNA, or misc_feature. A custom configuration file further allows modifying the colour of a gene and/or the name of the gene product to be displayed in the map (17,18). In the “Genes and Features” box (IIb), the user can select/deselect gene or feature classes (standard mode) or single genes (transcript mode), to be displayed or hidden in the map. In the box underneath (box IIc), detection methods for the inverted repeat (IR) regions present in most chloroplast genomes can be chosen. The last box (IId) of column II allows selection of restriction enzymes whose recognition sites are to be displayed in the map (allowing generation of a combined physical and restriction map). In column III, “Map Options” (box IIIa) and “Output Options” (box IIIb) can be specified. “Map Options” are, for example, inclusion of a graph of the GC content (available for circular maps) or the possibility to zoom into a specific region of the organellar genome, an option available for linear maps. In the transcript mode, the colour code for up- and down-regulated transcripts can be adjusted. With the “Output Options”, the user can choose between various kinds of vector and bitmap output file formats, and also specify the resolution of bitmap files. The “Action” box (IIIc) submits and resets jobs, but also allows loading example jobs that include demonstration files that can be downloaded for inspection by the user. In the “Results” box (IIId), the output files such as the physical map (Output-Graph) can be download individually by clicking on the respective symbol. By clicking the floppy disk symbol, all result and input files can be conveniently downloaded as a single zip archive.

**Figure 1.**
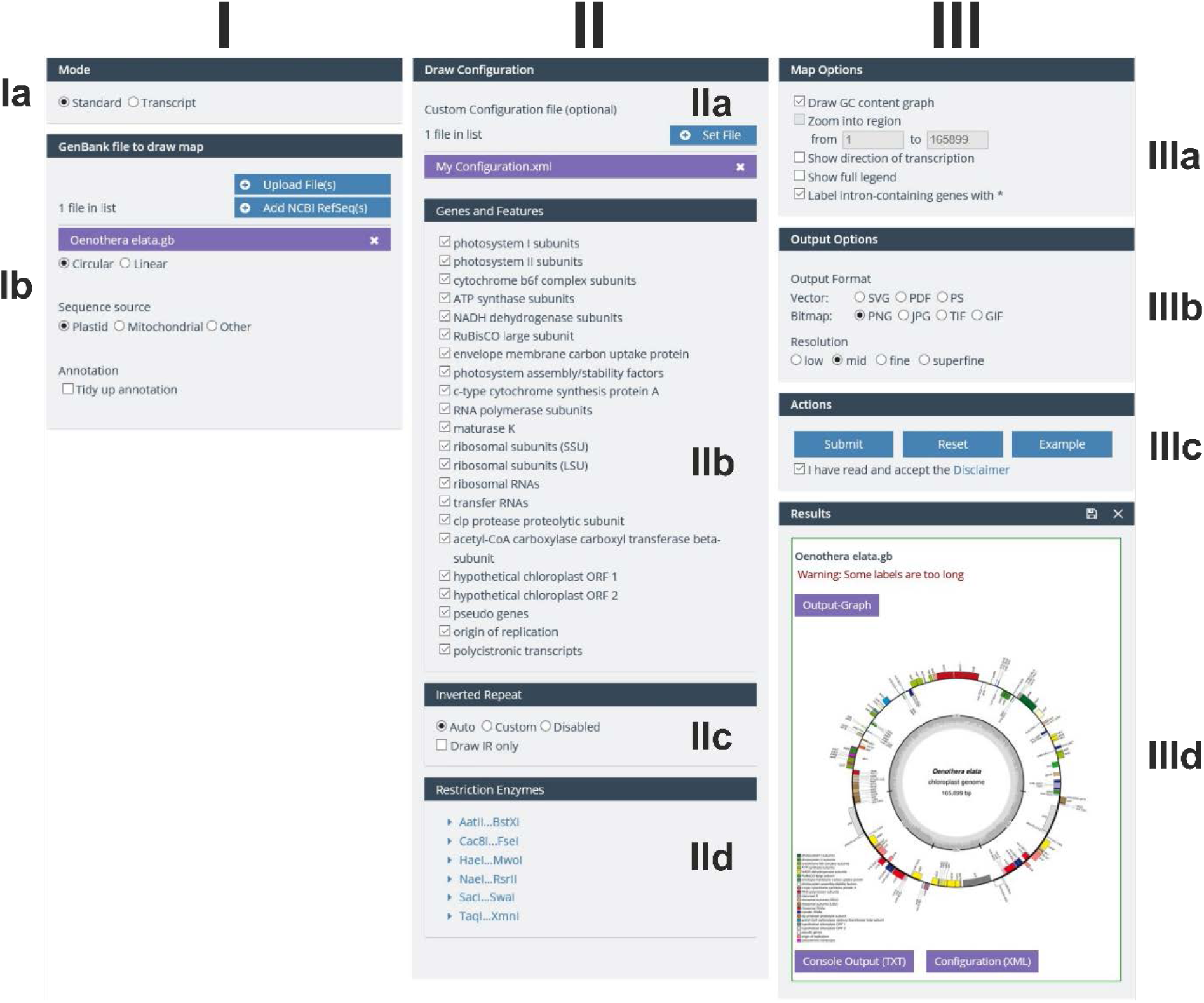
OGDRAW GUI in the standard mode. For details, see text.

**Figure 2.**
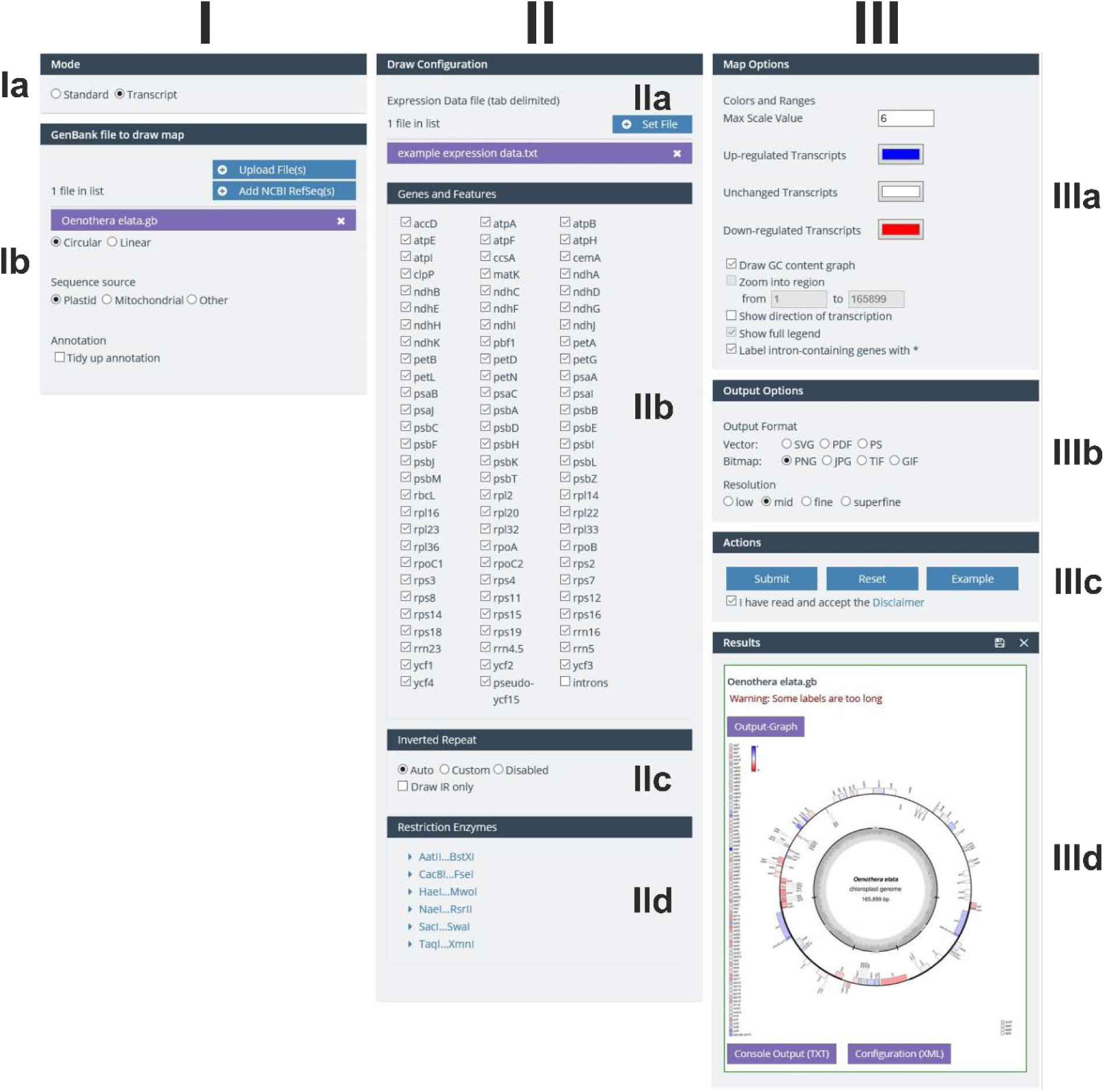
OGDRAW GUI in the transcript mode. For details, see text.

### The new features

With respect to the previous version v1.2 (18), OGDRAW v1.3.1 includes several new features:

(i) OGDRAW v1.3.1 has access to CHLOROBOX’s local copy of the organelle genome records of the NCBI RefSeq project (http://www.ncbi.nlm.nih.gov/genome/organelle/). The records are regularly updated, visualized as a phylogenetic tree, and their LOCUS, SOURCE and ORGANISM information is searchable by free text (15). Possible search queries are names of taxonomic ranks, scientific and (if present in the NCBI recorded) common names, the NCBI RefSeq accession number, or combinations thereof.
(ii) On the new OGDRAW server, the user can upload and process several individual as well as multi-GenBank files. In combination with selection of the GenBank files from the local NCBI RefSeq database, the large numbers of available organelle genomes for entire taxonomic clades can be visualized easily and very quickly. These large datasets can be downloaded as a single zip file (see above).
(iii) As new output option, modern vector graphic formats such as SVG and PDF were implemented.
(iv) OGDRAW v1.3.1 in its default parameters follows the convention to label intron-containing genes in organelle genomes with an asterisk (*). In the “Map Option” box, this optional feature can be deselected, however. If the user then selects “intron” in the “Genes and Features” box, the default parameters of the previous OGDRAW version are applied in that the intron is directly drawn into the gene as an empty box (Figure 3).
(v) In agreement with current annotation practise, operons can be displayed in the map as polycistronic transcription units, if annotated in the GenBank entry with the feature keys “prim_transcript” (17) and (as a newly included feature key) “operon”.
(vi) The D-loop of metazoan mitochondrial genomes is now drawn by default (Figure 4).
(vii) Genes or features that span start and end of submitted linear sequences are now displayed correctly by OGDRAW. While this feature is of minor relevance to the visualization of finalized annotations of organellar genomes (but also see Figure 4), its implementation became necessary to improve communication between OGDRAW and our organelle genome annotation pipeline GeSeq (15).

**Figure 3.**
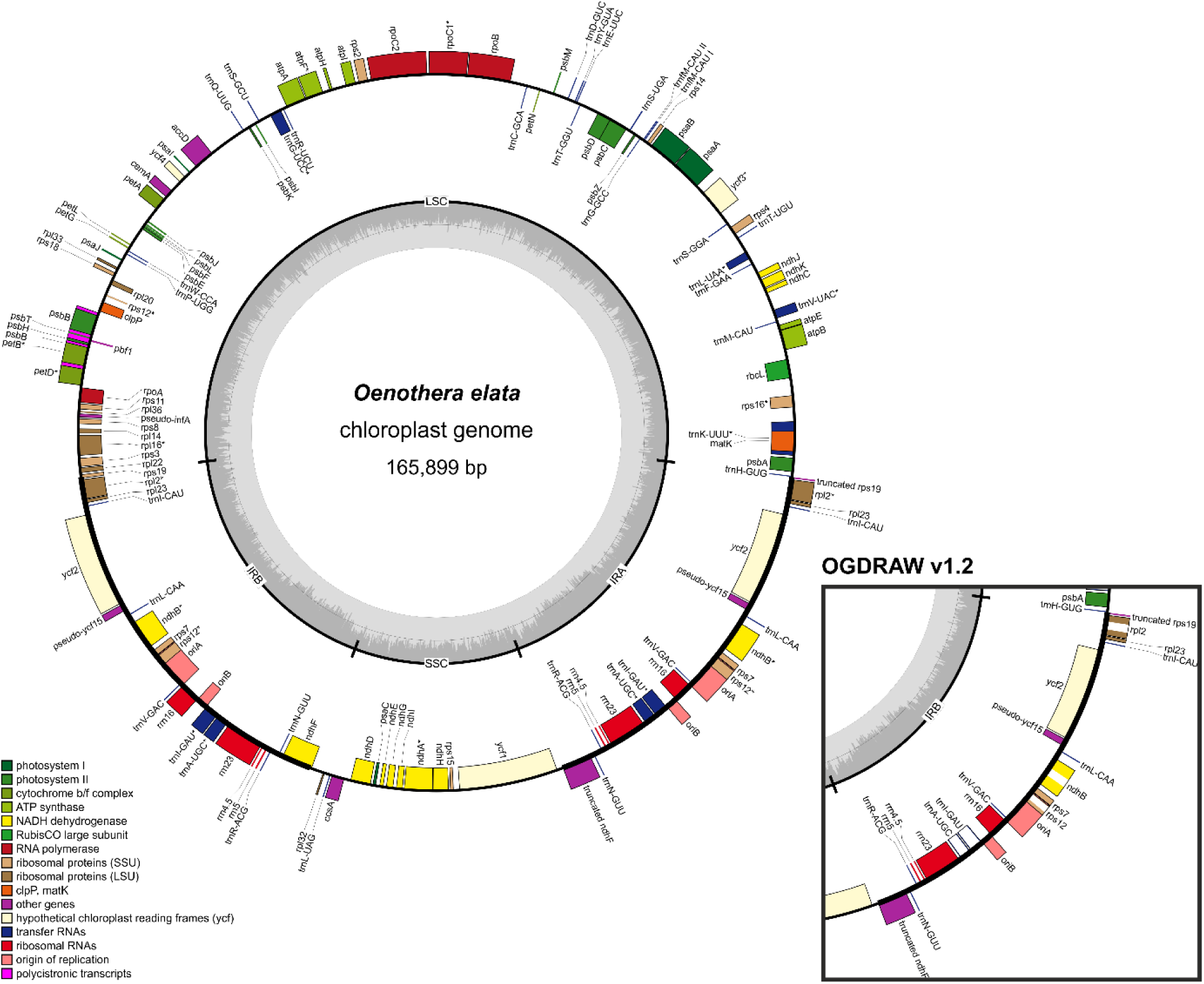
OGDRAW v1.3.1 output with default parameters illustrating the chloroplast genome of *Oenothera elata* (AJ271079.4). Genes inside the circle are transcribed clockwise, genes outside the circle counter clockwise. The circle inside the GC content graph marks the 50% threshold. Note that in contrast to earlier versions of OGDRAW (right box), intron containing genes are now marked by an asterisk (*) and introns are no longer directly drawn into the genes. Please also note that the inverted repeat A (IRA) is now designated as the right one on the map. For details see text.

**Figure 4.**
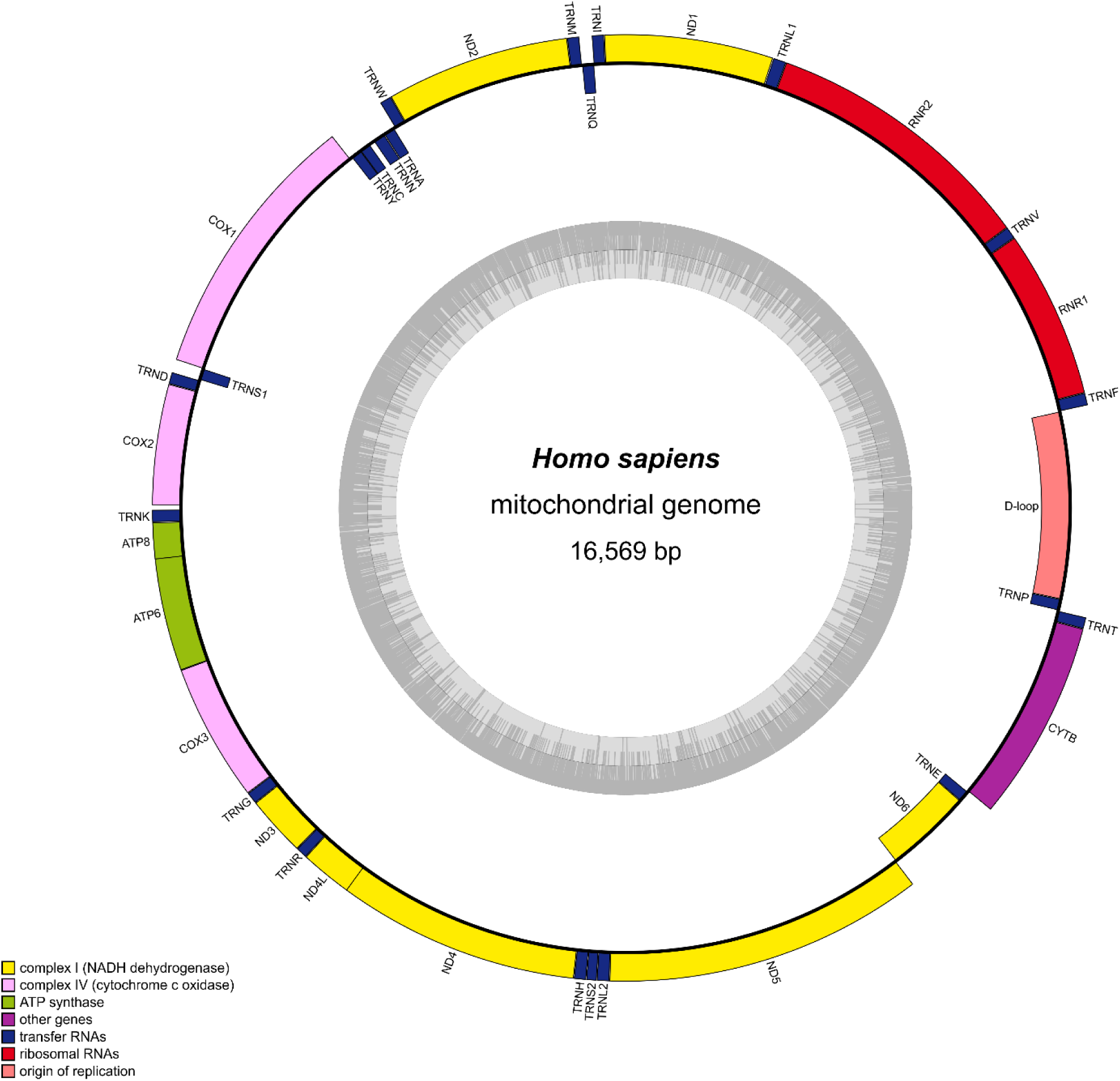
OGDRAW v1.3.1 output with default parameters illustrating the human mitochondrial genome (NC_012920.1). Genes inside the circle are transcribed clockwise, genes outside the circle counter clockwise. The circle inside the GC content graph marks the 50% threshold. Note that the D-loop, spanning start and end of the submitted linear sequence, is now drawn by the default.

### Bug fixes

A number of bugs were fixed in OGDRAW v1.3.1 and the most important ones are listed below:

(i) We revised the calculation of the stretch factor of linear maps. Previously, this feature did not work for many GenBank files.
(ii) OGDRAW now accepts GenBank files that contain an N or other IUPAC characters that are different from the four standard nucleotides A, T, G and C (19). Earlier versions of OGDRAW produced incorrect maps from such sequences.
(iii) With respect to v1.2, the new version of OGDRAW transposes inverted repeats A and B in chloroplast genomes (IRA and IRB; Figure 3). By default, IRA is now designated as the right repeat in the map, since nucleotide number 1 (set at approximately 3 o’clock by OGDRAW) is usually annotated as the first base of the large single copy region (LSC) flanked by IRA (20,21).

## Conclusion

For more than a decade, OGDRAW has provided the community with a user-friendly application to draw maps of organellar genomes. The program has become the standard in the field. The new version presented here (OGDRAW v1.3.1) provides improved functionality and versatility, and further increases user friendliness.

## Acknowledgement

We wish to thank the IT Service Team of the Max Planck Institute of Molecular Plant Physiology for excellent technical assistance.

## References

1. Tonti-Filippini, J., Nevill, P.G., Dixon, K. and Small, I. (2017) What can we do with 1000 plastid genomes? Plant J., 90, 808–818. https://doi.org/10.1111/tpj.13491 https://www.ncbi.nlm.nih.gov/pubmed/28112435

2. Wicke, S. and Schneeweiss, G.M. (2015) In Hörandl, E. and Appelhans, M. S. (eds.), *Next-Generation Sequencing in Plant Systematics*. Koeltz Scientific, Königstein, Germany, pp. 9–50. https://www.koeltz.com/product.aspx?pid=209039

3. Smith, D.R. (2016) The past, present and future of mitochondrial genomics: have we sequenced enough mtDNAs? Brief. Funct. Genomics, 15, 47–54. https://doi.org/10.1093/bfgp/elv027 https://www.ncbi.nlm.nih.gov/pmc/articles/PMC4812591/

4. Jin, J.-J., Yu, W.-B., Yang, J.-B., Song, Y., Yi, T.-S. and Li, D.-Z. (2018) GetOrganelle: a simple and fast pipeline for de novo assembly of a complete circular chloroplast genome using genome skimming data. bioRxiv, 256479. https://doi.org/10.1101/256479

5. Bakker, F.T., Lei, D., Yu, J., Mohammadin, S., Wei, Z., van de Kerke, S., Gravendeel, B., Nieuwenhuis, M., Staats, M., Alquezar-Planas, D.E. et al. (2016) Herbarium genomics: plastome sequence assembly from a range of herbarium specimens using an Iterative Organelle Genome Assembly pipeline. Biol. J. Linn. Soc., 117, 33–43. https://doi.org/10.1111/bij.12642

6. Jung, J., Kim, J.I., Jeong, Y.-S. and Yi, G. (2018) AGORA : Organellar genome annotation from the amino acid and nucleotide references. Bioinformatics, 34, 2661–2663. https://doi.org/10.1093/bioinformatics/bty196 https://www.ncbi.nlm.nih.gov/pubmed/29617954

7. Liu, C., Shi, L., Zhu, Y., Chen, H., Zhang, J., Lin, X. and Guan, X. (2012) CpGAVAS, an integrated web server for the annotation, visualization, analysis, and GenBank submission of completely sequenced chloroplast genome sequences. BMC Genomics, 13, 715. https://doi.org/10.1186/1471-2164-13-715 https://www.ncbi.nlm.nih.gov/pmc/articles/PMC3543216/

8. Wyman, S.K., Jansen, R.K. and Boore, J.L. (2004) Automatic annotation of organellar genomes with DOGMA. Bioinformatics, 20, 3252–3255. https://doi.org/10.1093/bioinformatics/bth352 https://www.ncbi.nlm.nih.gov/pubmed/15180927

9. Iwasaki, W., Fukunaga, T., Isagozawa, R., Yamada, K., Maeda, Y., Satoh, T.P., Sado, T., Mabuchi, K., Takeshima, H., Miya, M. et al. (2013) MitoFish and MitoAnnotator: a mitochondrial genome database of fish with an accurate and automatic annotation pipeline. Mol. Biol. Evol., 30, 2531–2540. https://doi.org/10.1093/molbev/mst141 https://www.ncbi.nlm.nih.gov/pmc/articles/PMC3808866/

10. Alverson, A.J., Wei, X., Rice, D.W., Stern, D.B., Barry, K. and Palmer, J.D. (2010) Insights into the evolution of mitochondrial genome size from complete sequences of *Citrullus lanatus* and *Cucurbita pepo* (Cucurbitaceae). Mol. Biol. Evol., 27, 1436–1448. https://doi.org/10.1093/molbev/msq029 https://www.ncbi.nlm.nih.gov/pmc/articles/PMC2877997/

11. Bernt, M., Donath, A., Jühling, F., Externbrink, F., Florentz, C., Fritzsch, G., Pütz, J., Middendorf, M. and Stadler, P.F. (2013) MITOS: Improved de novo metazoan mitochondrial genome annotation. Mol. Phylogen. Evol., 69, 313–319. https://doi.org/10.1016/j.ympev.2012.08.023 https://www.ncbi.nlm.nih.gov/pubmed/22982435

12. Sheffield, N.C., Hiatt, K.D., Valentine, M.C., Song, H. and Whiting, M.F. (2010) Mitochondrial genomics in *Orthoptera* using MOSAS. Mitochondrial DNA, 21, 87–104. https://doi.org/10.3109/19401736.2010.500812 https://www.ncbi.nlm.nih.gov/pubmed/20795780

13. Huang, D.I. and Cronk, Q.C.B. (2015) Plann: A command-line application for annotating plastome sequences. Appl. Plant Sci., 3, 1500026. https://doi.org/10.3732/apps.1500026 https://www.ncbi.nlm.nih.gov/pmc/articles/PMC4542940/

14. McKain, M.R., Hartsock, R.H., Wohl, M.M. and Kellogg, E.A. (2017) Verdant: automated annotation, alignment and phylogenetic analysis of whole chloroplast genomes. Bioinformatics, 33, 130–132. https://doi.org/10.1093/bioinformatics/btw583 https://www.ncbi.nlm.nih.gov/pmc/articles/PMC5408774/

15. Tillich, M., Lehwark, P., Ulbricht-Jones, E.S., Fischer, A., Pellizzer, T., Bock, R. and Greiner, S. (2017) GeSeq-versatile and accurate annotation of organelle genomes. Nucleic Acids Res., 45, W6–W11. https://doi.org/10.1093/nar/gkx391 https://www.ncbi.nlm.nih.gov/pmc/articles/PMC5570176/

16. Gissi, C., Iannelli, F. and Pesole, G. (2008) Evolution of the mitochondrial genome of Metazoa as exemplified by comparison of congeneric species. Heredity, 101, 301–320. https://www.nature.com/articles/hdy200862 https://www.ncbi.nlm.nih.gov/pubmed/18612321

17. Lohse, M., Drechsel, O. and Bock, R. (2007) OrganellarGenomeDRAW (OGDRAW): a tool for the easy generation of high-quality custom graphical maps of plastid and mitochondrial genomes. Curr. Genet., 52, 267–274. https://doi.org/10.1007/s00294-007-0161-y https://www.ncbi.nlm.nih.gov/pubmed/17957369

18. Lohse, M., Drechsel, O., Kahlau, S. and Bock, R. (2013) OrganellarGenomeDRAW-a suite of tools for generating physical maps of plastid and mitochondrial genomes and visualizing expression data sets. Nucleic Acids Res., 41, W575–W581. https://doi.org/10.1093/nar/gkt289 https://www.ncbi.nlm.nih.gov/pmc/articles/PMC3692101/

19. Cornish-Bowden, A. (1985) Nomenclature for incompletely specified bases in nucleic acid sequences: recommendations 1984. Nucleic Acids Res., 13, 3021–3030. https://doi.org/10.1093/nar/13.9.3021 https://www.ncbi.nlm.nih.gov/pmc/articles/PMC341218/

20. Shinozaki, K., Ohme, M., Tanaka, M., Wakasugi, T., Hayashida, N., Matsubayashi, T., Zaita, N., Chunwongse, J., Obokata, J., Yamaguchi-Shinozaki, K. et al. (1986) The complete nucleotide sequence of the tobacco chloroplast genome: its gene organization and expression. EMBO J., 5, 2043–2049. https://doi.org/10.1002/j.1460-2075.1986.tb04464.x https://www.ncbi.nlm.nih.gov/pmc/articles/PMC1167080/

21. Ohyama, K., Fukuzawa, H., Kohchi, T., Shirai, H., Sano, T., Sano, S., Umesono, K., Shiki, Y., Takeuchi, M., Chang, Z. et al. (1986) Chloroplast gene organization deduced from complete sequence of liverwort *Marchantia polymorpha* chloroplast DNA. Nature, 322, 572–574 https://doi.org/10.1038/322572a0

